# Learning of goal-relevant and irrelevant complex visual sequences in human V1

**DOI:** 10.1101/261255

**Authors:** Clive R. Rosenthal, Indira Mallik, Cesar Caballero-Gaudes, Martin I. Sereno, David Soto

## Abstract

Learning and memory are supported by a network involving the medial temporal lobe and linked neocortical regions. Emerging evidence indicates that primary visual cortex (i.e., V1) may contribute to recognition memory, but this has been tested only with a single visuospatial sequence as the target memorandum. The present study used functional magnetic resonance imaging to investigate whether human V1 can support the learning of multiple, concurrent, and complex visual sequences involving discontinous (second-order) associations. Two peripheral, goal-irrelevant but structured sequences of orientated gratings appeared simultaneously in fixed locations of the right and left visual fields alongside a central, goal-relevant sequence that was in the focus of spatial attention. Pseudorandom sequences were introduced at multiple intervals during the presentation of the three structured visual sequences to provide an online measure of sequence-specific knowledge at each retinotopic location. We found that a network involving the precuneus and V1 was involved in learning the structured sequence presented at central fixation, whereas right V1 was modulated by repeated exposure to the concurrent structured sequence presented in the left visual field. The same result was not found in left V1. These results indicate for the first time that human V1 can support the learning of multiple concurrent sequences involving complex discontinuous inter-item associations, even peripheral sequences that are goal-irrelevant.

## 1 Introduction

Primary visual cortex (V1) is typically thought to carry out primitive low-level visual computations and perceptual learning of low-level visual features (Sasaki, Nanez, & Watanabe, 2010), but there is now emerging evidence that V1 can play a role in higher-order functions such as the learning and recognition of visuo-spatial sequences in humans (Rosenthal, Andrews, Antoniades, Kennard, & Soto, 2016), and in model organisms (Cooke, Komorowski, Kaplan, Gavornik, & Bear, 2015; Gavornik & Bear, 2014). Functional coupling between V1 and putative memory substrates has been identified during non-conscious recognition (Rosenthal et al., 2016), incidental statistical learning of visible items (Turk-Browne, Scholl, Johnson, & Chun, 2010), conscious recall of tone-grating pairs (Bosch, Jehee, Fernández, & Doeller, 2014) and cue-action associations (Hindy, Ng, & Turk-Browne, 2016). However, the factors that modulate V1 activity during the learning of complex memoranda remain to be systematically evaluated.

The aim of the present functional magnetic resonance imaging (fMRI) study was to assess whether human V1 can support concurrent learning and recognition memory of three complex visual sequences presented at goal-relevant (attended) locations and also at peripheral, goal-irrelevant regions of the visual field subtended by V1. Prior research has implicated V1 in learning and memory for single or simple visual sequences (Rosenthal et al., 2016; Cooke et al., 2015; Gavornik & Bear, 2014). For instance, a recent fMRI-study by Rosenthal et al. (2016) showed a role of V1 in learning a single, goal-relevant non-conscious second-order conditional visuospatial sequence of targets specified at four locations. This study, however, could not address whether multiple concurrent, goal-relevant and goal-irrelevant sequences can be encoded in V1. Also, it is also unknown whether or not higher-order learning effects for goal-irrelevant sequences occur in retinotopic V1 areas. The present study tackled these questions.

Our approach to understand perceptual sequence learning of goal-relevant and -irrelevant sequences was to assess changes in hemodynamic responses as a proxy for sequence-specific learning and also offline behavioural tests of recognition memory performance for the trained sequences. Importantly, however, any behavioural measures collected in a task context in which the previously irrelevant peripheral sequences are made goal-relevant may yield only noisy estimates of newly acquired knowledge due to diminished transfer-appropriate processing or violation of the encoding specificity (Tulving & Thomson, 1973; Morris, Bransford, & Franks, 1977; Mulligan & Lozito, 2006). Hence although we would not necessarily expect successful recognition memory for the peripheral goal-irrelevant sequences, the hemodynamic response can however provide an indirect yet dynamic measure of newly acquired sequence knowledge as it unfolds during learning (Karuza et al., 2013; McNealy, Mazziotta, & Dapretto, 2006; Rugg et al., 1998).

We thus assessed the brain responses in V1 associated with the repeated presentation of a visual sequence presented at fixation and also tested whether right and left V1 exhibited learning-related activity associated with the repeated presentation of the peripheral, goal-irrelevant, but structured, higher-order visual sequences in its corresponding contralateral visual receptive fields. Evidence of V1 modulation in either peripheral irrelevant location would be consistent with a model of V1 at the interface between perception and memory functions, which can operate independently of goal-directed strategic control factors.

## 2 Methods

### 2.1 Participants

Seventeen undergraduate students were recruited (mean age of 22 years; nine female). All participants gave informed written consent to take part in accordance with the terms of approval granted by the local research ethics committee and the principles expressed in the Declaration of Helsinki, and were naive to the purpose of the experiment. All participants reported normal or corrected-to-normal vision, and had no history of neurological disease.

### 2.2 Experimental procedure

Participants underwent fMRI scanning as they viewed a visual display on which three different visual sequences were presented. The sequence presented at the center of the screen involved changes in the color of a circle (white, light grey, dark grey and black). The sequences presented in the left and right visual fields involved changes in the orientation of a grating (i.e., 0, 45, 90 or 135 degrees from the vertical). Each sequence was comprised of 12 stimuli following a second-order conditional (SOC) rule (Reed & Johnson, 1994), whereby at the lowest structural level the current stimulus is contingent on the two preceding stimulus. The sequences used were as follows: SOC1: 3 4 2 3 1 2 1 4 3 2 4 1; SOC2: 3 4 1 2 4 3 1 4 2 1 3 2; SOC3: 2 4 2 1 3 4 1 2 3 1 4 3, where 1 2 3 4 represents a stimulus type). For instance, SOC1 comprises the following twelve deterministic SOC triplets (i.e., the lowest structural unit): 342 423 231 312 121 214 143 432 324 241 413 134 - accordingly, the SOC context (dependencies/determinacies) for 2 in the 3rd location is 3,4; for 3 in the 4th location, the SOC context it is 4,2; for 2 in the 6th location, the SOC context is 3,1; for 3 in the 9th location, it is 1,4; and, for 2 in the 10th location, the SOC context is 4,3. Each of the SOC sequences appeared at a fixed location across subjects (left: SOC1; center: SOC2, and right: SOC3). Note that all sequences were equated in terms of all salient features that are related to ease with which each sequence could be learned: namely, simple frequency of positions, first-order transition frequency, reversal frequency, rate of full coverage, and the number of second-order conditional triplets.

Participants were instructed to discriminate the current color of the central sequence by pressing one of four buttons. The central sequence was goal-relevant and hence in the focus of attention. The peripheral sequences were designated goal-irrelevant by instructing participant to ignore the changes in orientations of the gratings and only respond to the colour changes in the central sequence and not to respond to the orientations of these peripheral sequences. The phase of the gratings was varied independently of the sequences of orientations, and varied randomly across trials to preclude neural adaptation effects of the phase dimension. The three SOC sequences at each of the three locations were presented on a loop, but were asynchronously interspersed with the presentation of the 12 element pseudorandom sequences. The serial order of the items was pseudorandom, such that the same item did not appear consecutively (i.e., there was no immediate repetitions of a colour or grating orientation) and items appeared with equal frequency for each orientation or colour. Each pseudorandom sequence was unique for each of the 3 runs and for each sequence at each location.

In line with prior studies in sequence learning, we elected to use a pseudorandom sequence, instead of a non-trained second-order conditional sequence on baseline blocks to facilitate learning of the three target SOC sequences, as was the case in our prior study (Rosenthal et al., 2016). In particular, we have found that presentation of novel second-order conditional sequences as baseline blocks impairs the learning of the target structured 12-item second-order conditional sequence. Furthermore, many other notable human and infra-human studies of the brain bases of sequence learning have similarly contrasted the learning of visible second-order conditional sequences with pseudorandom baseline blocks (Schendan, Searl, Melrose, & Stern, 2003; Ergorul & Eichenbaum, 2006; Poldrack et al., 2005; Gheysen, Van Opstal, Roggeman, Van Waelvelde, & Fias, 2011).

Pseudorandom sequences were introduced at multiple intervals during the presentation of the three SOC sequences, as depicted in Figure 1. Pseudorandom sequences were always repeated 4 times, hence comprising 48 stimulus presentations or trials. Stimuli at all three locations were each presented for 1 second with an interval of 0.25 seconds between each stimulus. The pseudorandom sequence appeared twice per run at different times for each of the 3 sequences. The temporal position within a run of each of the pseudorandom sequences was consistent across participants. Figure 1 (right panel) illustrates the structure of each of the 3 training runs. There were 672 trials on each of the three training runs and each lasted for 14 minutes.

**Figure 1:**
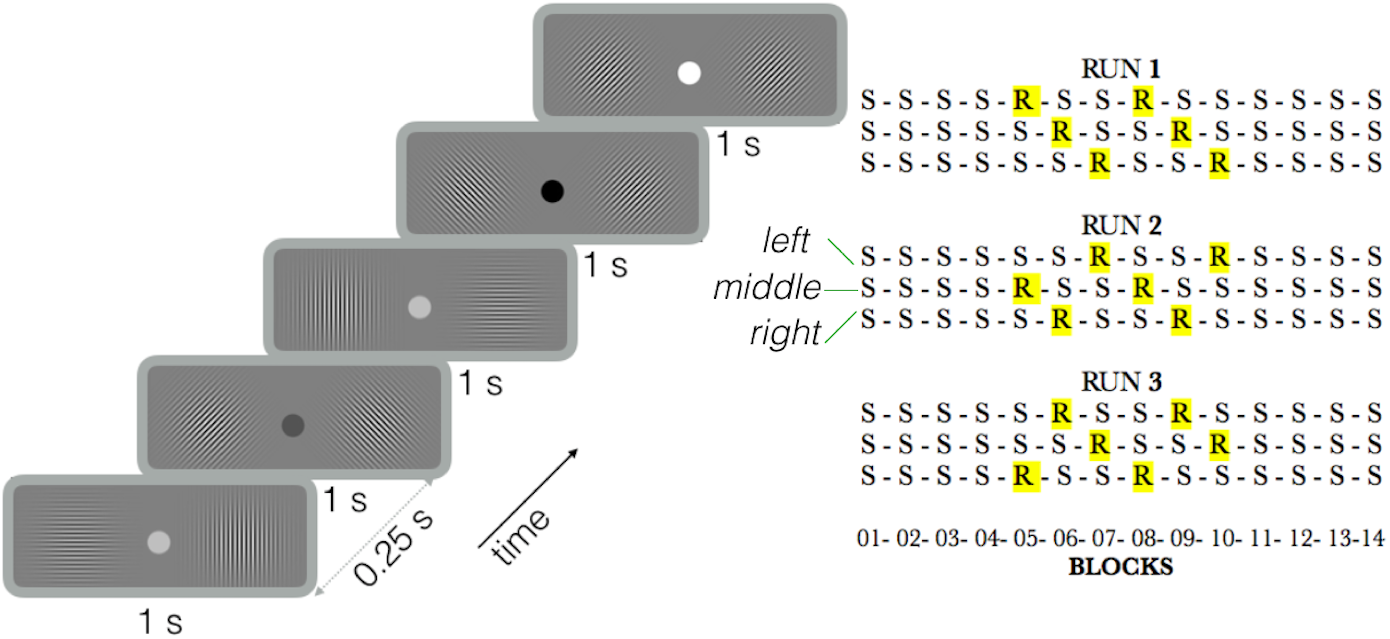
Illustration of the sequence training protocol inside the MRI scanner. The right panel indicates the structure of each of the 3 training runs (S: for the visual second-order conditional sequence; R: for pseudorandom blocks, comprised of a 48 trials each). The top row corresponds to the left sequence, the middle to the central sequence and the bottom row within each run corresponds to the right sequence. Note that each of the pseudorandom sequences were presented asynchronously across the three sequences and scanning runs.

### 2.3 Recognition test outside of the scanner

After training, the participants performed three recognition tests outside of the scanner, one for each of the three 12-element second-order conditional sequences presented at central fixation and left/right positions of the visual field. The order of the tests was counterbalanced across participants. On each trial of the recognition test, participants were shown either an old or a new sequence which served as a recognition cue (6-element sequences). Twelve 6-element sequences were generated from each SOC. Each trial 6-element sequence on the recognition tests was generated by starting from a different ordinal position of the 12-element SOC sequence for six consecutive locations.

Old sequences corresponded structured sequences presented during the training phase inside the scanner. As an example, the left SOC sequence during training was presented in the left visual field during the recognition memory test and the cues based on the trained ’old’ sequences were interspersed with new sequences. New sequences were also based on a SOC rule, but had not been presented to the participant during the training phase. The participant was asked to indicate whether each of the sequences presented on each trial was ’old’ or ’new’. Then, participants indicated the confidence in their response (from 1-guess; 2-fairly confident; 3-sure). Participants were allowed to simulate the motor responses given during training when viewing the central sequences during the recognition test.

Receiver-operating-characteristic (ROC) analyses were used to derive measures of type-1 sensitivity (i.e., ability to distinguish old/new sequences) and type-2 sensitivity (i.e., how confidence relates to memory accuracy). For the type-1 analyses, the ’signal’ and ’noise’ were defined as old and new retrieval cues, respectively. A ’hit’ was, therefore, a correct response (’old’) to an old trained (six-element sequence) retrieval cue and a ’correct rejection’ was a correct response (’new’) to a new untrained (six-element sequence) retrieval cue. A ’false alarm’ was an incorrect response (’old’) to a new retrieval cue and a ’miss’ was an incorrect response (’new’) to an old retrieval. To obtain the type-1 ROC curves, we plotted the probability of hits as a function of the probability of false alarms for all possible decision criteria. The different points in the ROC curve were obtained by calculating cumulative probabilities for hits and false alarms along a recoded confidence continuum ranging from 3 (i.e., certain) to 1 (i.e., least certain) in signal present trials (old retrieval cues), and from 3 (i.e., certain) to 1 (i.e., least certain) in noise trials (new retrieval cues). On the basis of simple geometry, we computed the area under the ROC curve as a distribution-free measure of the discriminability. For the type-2 sensitivity analyses, the area under the ROC curve was calculated by plotting the cumulative probability of correct responses (either hit or correct rejection) and the cumulative probability of incorrect responses (either false alarm or miss) across the different levels of confidence. The area under the ROC curve was estimated as distribution-free measure of metacognitive ability (Fleming, Weil, Nagy, Dolan, & Rees, 2010; Kornbrot, 2006). Data processing and analyses were performed in R (version 3.2.2).

### 2.4 MRI data acquisition

MRI scanning was performed in a Siemens Avanto 1.5 T MRI scanner using a receive-only 32-channel head coil and body transmit. A screen was mounted at the end of the scanner bore and visual stimuli were projected onto it from the console room through a wave guide. Participants viewed the screen using a 45 deg mirror. Functional volumes for both the training phase, functional localiser, and retinotopy consisted of multi-slice T2*-weighted echoplanar images (EPI) with blood oxygenation level dependent (BOLD) contrast with a multiband acceleration factor of 4. We used the following scanning parameters to achieve whole brain coverage: TR=1000 ms, TE=54.8 ms, 40 coronal slices, 3.2 mm slice thickness, no interslice gap, and FoV=205×205 mm (3.2 × 3.2 mm inplane voxels). There were 844 whole-brain scans per training run. To facilitate anatomical localization and cross-participant alignment, a high-resolution 1×1×1 mm whole-brain structural T1-weighted, magnetization prepared rapid gradient echo (MP-RAGE) scan was acquired for each participant (FOV=256×224 mm, 176 partitions, TR=8.4 ms, TE=3.57 ms, TI=1000 ms, inversion spacing=2730 ms, BW=190 Hz/pixel).

### 2.5 Functional MRI preprocessing

We used FEAT (fMRI Expert Analysis Tool) Version 6.0, as part of FSL (www.fmrib.ox.ac.uk/fsl). The first 10 EPI volumes were removed to account for T1 equilibrium effects. Non-brain removal was performed using Brain Extraction Tool. Volume realigment of functional scans was carried out using FMRIB’s Linear Image Registration Tool MCFLIRT. Scans were realigned relative to the middle scan. We applied a 50 s high-pass temporal filtering to remove low frequency noise, and spatial smoothing using a FWHM Gaussian kernel of 6 mm.

### 2.6 Retinotopic mapping

Retinotopic mapping was performed to identify right and left human V1 in each individual. We used standard phase-encoded polar angle retinotopic mapping (Sereno et al., 1995). Participants fixated centrally while viewing a clockwise or counterclockwise rotating pie-shaped ’wedge’ with a polar eccentricity. The duration of the scan was 8 min 32 secs (512 volumes). Fourier methods were applied to obtain polar angle maps of the cortex and delineate the borders of right and left V1.

### 2.7 Functional localizer

A functional localiser scan was conducted in all participants and involved the presentation of items that were flashed at the same three different locations. A block design was used. At each location, there were 24 presentations of gratings (250 ms each with a 250 ms inter-grating interval) in pseudorandom orientations. Following each block, there was a rest period of 12 seconds. The phase of the gratings also varied randomly across trials. We obtained contrast of parameter estimates of brain responses to the spatial position of the peripheral sequences (left>right and right>left positions).

The parameter estimates from the localiser were used to define another mask comprising the voxels in the V1 retinotopic masks that responded to the position of the gratings during the training phase (e.g., define the voxels within the right V1 mask that showed a response to the left-side position of the gratings and likewise for left V1). This area in V1 was delineated by finding the intersection between the V1 retinotopy and the functional localiser maps using fslmaths -mas tool. Hence, we isolated the area of V1 that was most selective of the location of the right and left sequences. We initially derived masks comprising the 10 most responsive voxels and repeated the procedure considering the 5, 20 and 40 most responsive voxels to further assess the reliability of the results.

### 2.8 MRI Statistical Analyses

Time-series statistical analyses were conducted using FILM (FMRIB’s Improved Linear Model) with local autocorrelation correction. The data were analyzed using voxelwise time series analysis within the framework of the general linear model. A design matrix was generated in which each individual presentation of a grating was modelled with a double-gamma hemodynamic response function. Explanatory regressors were created for each of the structured and pseudorandom sequences. Standard motion realignment parameters were included in each individual subject’s general linear model as nuisance regressors. We included further nuisance regressors for those volumes corresponding to motion outliers using the FSL motion outliers function. The root mean squared (RSM) head position difference to the reference volume was used as metric to detect fMRI timepoints corrupted by large motion. The threshold was set to the FSL default, namely, the 75 percentile + 1.5 inter quartile range of the distribution of RSM values of each run. A confound matrix was generated and used in the GLM to completely remove the effects of these timepoints on the analysis. This is intended to deal with the effects of intermediate to large motions, which corrupt images beyond anything that the linear motion parameter regression methods can fix.

Sequence blocks were modeled in chunks of 48 trials in order to contrast activity between structured and pseudorandom baseline blocks (S and R in Figure 1); 48 trials were selected as a multiple of 12-element unit of SOC sequence that allowed the number of trials on structured sequence blocks to be equated with the pseudorandom blocks, as per our prior study (Rosenthal et al., 2016).

For the analysis of the central sequence, we first derived contrast of parameter estimates between the structured relative to the pseudorandom sequences (S<R) within each run. S<R contrast of parameter estimates were obtained as follows. First, we compared the first central pseudorandom sequence block in a run with the preceding structured central sequences that were not flanked by random sequences in the right and left visual fields (e.g., see Figure 1). Therefore, the two structured blocks preceding the first central pseudorandom were not considered in the S<R contrast. Instead, we selected the preceding structured block (i.e., the third one preceding the first central pseudorandom sequence, see Figure 1). This was done to exclude the possibility that BOLD responses to the critical central structured sequence were counfounded by hemodynamic responses changes from the peripheral pseudorandom sequence presented concurrently. Secondly, we also derived contrasts of parameter estimates by comparing the structured block of 48 trials that immediately preceded the first pseudorandom block on each of the three runs. The results obtained from these two analysis strategies were similar.

Following the computation of within-run contrast of parameter estimates (S<R), we performed across-run within-subject (fixed-effects) analyses using lower level parameter estimates to test for a linear training effects across the 3 runs (e.g., -1 0 1) and likewise exponential training effects (e.g., -2 -0.5 2.5). Finally, the parameter estimates from these linear and exponential contrasts across the three learning runs were converted to standard MNI 2 mm template with linear transformation and passed to a higher-level across subjects one-sample t-test to assess brain regions consistent across participants that were associated with increased training effects in the structured relative to the pseudorandom sequence using FLAME (Local Analysis of Mixed Effect). Statistical maps were thresholded using clusters determined by Z > 2.3 and a corrected cluster extent significance threshold of p=0.05, using Gaussian Random Field Theory. Across participants one-sample t-tests were also performed separately on each of the 3 runs to assess S<R brain actitivy differences on each run. These considerations applied also to the psychophysiological interaction (PPI) analyses described below.

We also used non-parametric permutation-based one-sample t-tests to further assess the statistical significance of the results. The FSL Randomise program was used (Winkler, Ridgway, Webster, Smith, & Nichols, 2014) with threshold-free cluster enhancement (TFCE). TFCE is a method that enhances values within cluster-like structures, while preserving voxel-level inference (Smith & Nichols, 2009). We performd 6 mm smoothing of the variance which is typically included to increase study power when sample sizes are lower than 20 in non-parametric testing (Nichols & Holmes, 2002).

For the analysis of the peripheral right and left sequence, we adopted the following approach. Parameter estimates were derived in native functional space for each structured and pseudorandom sequence block (Figure 1) for each training run and participant. We then used the V1 masks obtained from the retinotopy and the functional localiser to extract the parameter estimates of V1 activity across the training phase, separately for the right and left V1 masks. For each run, we then performed paired-sample t-tests comparing V1 activity estimates in the first pseudorandom sequence and the inmediately preceding structured sequence.

Percent signal changes were calculated for visualization of the time course of the learning effects using the following formula (Mumford, 2007): [contrast image / (mean of run)] * scaling factor * 100, where the scaling factor = baseline-to-max range) /(contrast fix).

### 2.9 Psychophysiological interaction (PPI) analyses

A seed-voxel PPI based approach was used to identify signals of interest that would be missed in standard subtraction based analyses (O’Reilly, Woolrich, Behrens, Smith, & Johansen-Berg, 2012). A 6 mm radius posterior cingulate/precuneus mask was drawn and based on the peak voxels of the posterior cingulate/precuneus that showed effects of training. This mask was used to define the seed region’s time course for the PPI analyses. The aim was to assess whether the temporal correlation between posterior precuneus and primary visual cortex was modulated during the learning of the goal-relevant, second-order conditional sequence.

We then estimated a model for each participant that included the same psychological regressors as outlined above, for the onsets of each sequence, and, critically, for the PPI model, a physiological regressor for the time course of the region-of-interest, and psychological × physiological interaction regressors for the PPI. These new regressors were, therefore, added to the previous first-level model for each participant/run. Parameter estimates for structured versus pseudorandom conditions based on the PPI regressors were derived in a similar fashion to the analyses outlined above (i.e., using both fixed-effects analysis across runs followed by mixed-effect analysis across participants), which, here, directly compared the changes in functional coupling associated with training. Given our a priori interests in V1(Rosenthal et al., 2016), the higher-level analyses (across subjects) were performed using a region-of-interest approach, with a target occipital mask in standard space derived from the FSL Harvard-Oxford Atlas.

## 3 Results

### 3.1 Behavioural performance during training

We assessed the response time (RT) and the accuracy of manual responses to each item color of the central sequence as a function of the nature of the sequence (structured vs pseudorandom). Overall, the results indicate that the repeated exposure to the deterministic discontinuous associations of the structured sequences affected somatomotor performance relative to each novel pseudorandom sequence. We conducted an ANOVA on the manual RTs with run(1,2,3) and sequence type (S,R) as factors. We compared the first R sequence and the preceding S sequence of each run. There was no main effect of run on RTs (F(2,32)=1.95, p=0.159). The main effect of sequence type was significant (F(2,32)=11.67, p=0.004), with slower performance on R relative to S sequence; however this effect did not interact with run. (F(2,32=0.607, p=0.551). The accuracy data showed a consonant pattern with the RTs. There was no main effect of run on accuracy (F(2,32)=0.177, p=0.838). The main effect of sequence type was significant (F(2,32)=14.913, p=0.001), with impaired accuracy on R relative to S sequence blocks; however this effect did not interact with run (F(2,32=2.287, p=0.118). Figure 2 illustrates these results. Taken together, these results indicate that the statistical structure of the central goal-relevant sequence was learned.

**Figure 2:**
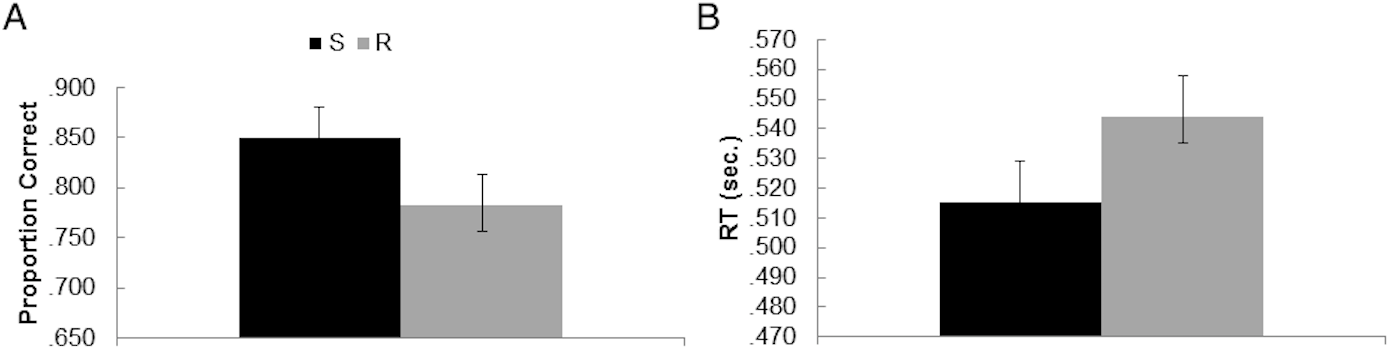
Behavioural results of the learning phase collapsed across runs (A) Mean proportion of correct responses in S and R blocks (B) Mean RTs in S and R conditions.

### 3.2 Recognition performance outside the scanner

Following the fMRI training task, a surprise recognition test was administered to test the memory of participants for each of the 3 sequences. To derive sensitivity measures of recognition performance we computed the area under the type-1 ROC for ’old’/’new’ discrimination. For the sequence presented at central fixation, memory sensitivity was marginally above chance (t(16)=1.882, p=0.078, mean=0.565). Type-2 sensitivity performance, namely, the extent to which confidence ratings predicted the accuracy of the recognition response showed a similar trend (t(16)=1.911, p=0.074, mean=0.544).

However, additional analyses of recognition performance based on memory confidence evinced that old and new probes were processed differently. Notably, participants were more confident in their responses when judging old sequences relative to new sequences (t(16)=2.2706, p=0.036, mean confidence old=1.926; mean new=1.784), indicating that confidence was diagnostic of the memory status (old/new) of the sequences. The memory sensitivity and confidence results are presented in Figure 3.

**Figure 3:**
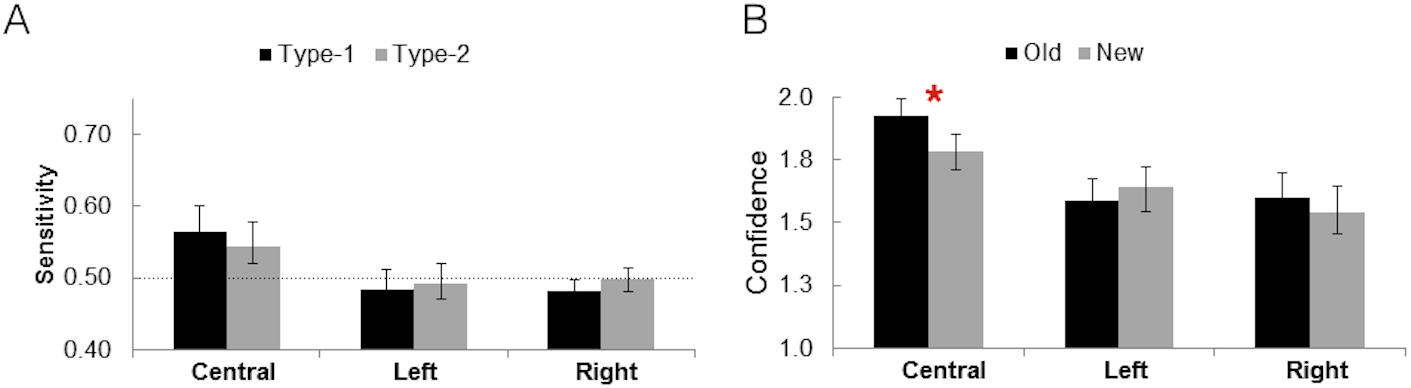
Behavioural results of the recognition test outside the scanner (A) Type-1 and type-2 memory sensitivity to old and new sequences (B) Memory confidence for old and new sequences.

Further results presented next indicated that participants were unable to recognise either the left or the right sequences.

Analyses of the recognition test performance for the left sequence showed that the area under the type-1 and type-2 ROC was no different from chance (t(16)=0.543, p=0.594, mean= 0.484 for type-1 ROC, and t(16)=0.352, p=0.729, mean=0.492 for type-2 ROC performance). Moreover, the results showed that participants were no more confident in responding to old vs. new left sequences (t(16)=1.0371, p=0.31, mean confidence old=1.593; mean new=1.647).

There was also no evidence of recognition memory for the right sequence. Type-1 and type-2 performance were no different from chance (t(16)=1.075, p=0.298, mean 0.483, and t(16)=0.101, p=0.92, mean=0.498). Participants confidence ratings did not discriminate between old and new left sequences (t(16)=1.165, p=0.261, mean confidence old=1.598; mean new=1.544).

Nonetheless, additional analyses conducted for the peripheral sequences showed differences in the latency of responses to old versus new recognition cues. Recognition reaction times (RTs) can provide a cogent metric of newly acquired sequence-specific knowledge because new six-element cues differed only in terms of study status, having been equated with the old six-element cues across all salient structural properties (Shanks, Channon, Wilkinson, & Curran, 2006). We computed a measure of recognition performance based on reaction time (RT) differences for recognition hits and correct rejections (Shanks et al., 2006). An ANOVA with sequence (left, right) and sequence status (old, new) was conducted on the mean RT differences of the correct responses for old and new probes. One participant had to be discarded because he had no correct responses to new items of the right sequences. There was a significant effect of the old/new status (F(1,15)=16.6, p=0.001) but no effect of the visual field of the sequence (F(1,15)=0.460, p=0.51) and no interaction between factors (F(1,15)=1.775, p=0.2). This indicates the presence of learning effects indexed by the recognition RTs for left (mean old RT = 1.108; mean new RT = 1.307) and right sequences (mean old RT = 1.059 s; mean new RT = 1.603 s).

### 3.3 fMRI Results

A key question of the present study is whether the V1 is involved in supporting the learning of complex deterministic discountinous associations in the structured peripheral goal-irrelevant sequences of oriented gratings.

For the analysis of the central sequence, we computed statistical contrasts of parameter estimates between structured (S) and pseudorandom (R) sequences (i.e., comparing S<R) on each of the three training runs (see Methods). This S<R contrast predicts increased response to the presentation of (novel) pseudorandom sequences relative to the repeated representation of the structured sequence and this contrast was motivated by our prior fMRI perceptual sequence learning study (Rosenthal et al., 2016), and is in keeping with a plethora of evidence for novelty effects in memory paradigms which are reflected by increases in hemodynamic responses (Ranganath & Ritchey, 2012).

First, BOLD responses to the first R central sequence were compared to a preceding structured sequence that was not flanked by pseudorandom sequences (see Methods) in order to minimise the concern that BOLD responses to the critical central structured sequence were counfounded by hemodynamic responses changes from the peripheral pseudorandom sequences presented concurrently. We then tested for linear and exponential effects of training on BOLD responses (e.g., increased differential S<R response in run 3 relative to run 1). Learning the structure of the repeated sequence ought to be reflected in brain activity changes across the training runs whenever a non-structured, pseudorandom sequence was presented.

The analyses showed significant clusters in the posterior cingulate gyrus (MNI 0 -48 16, Z=4.26) extending into the retrosplenial cortex, and the posterior precuneus in the the vicinity of intracalcarine sulcus, the ventromedial prefrontal cortex (MNI, 2 64 2, Z=3.41). All of these regions displayed significant linear and exponential changes in activity across runs (Z > 2.3, p < 0.05, whole brain corrected using cluster-based random-field theory with family wise error correction).

Secondly, we performed the same analyses by comparing the structured sequence block immediately preceding a pseudorandom sequence block at similar timepoint across the runs. The results obtained from this second analysis were similar to the first except that the ventromedial prefrontal cortex did not survive the threshold for statistical significance. Activity was restricted to the posterior cingulate and posterior precuneus.

To examine further the effects of training, we performed group-analyses separately for each of the three runs. On the basis of prior studies of learning SOC sequences (Schendan et al., 2003; Albouy et al., 2008), training effects were predicted to be most evident on the later third training run. Differences between the structured and the pseudorandom sequence blocks in run 1 and in run 2 did not survive the significance threshold. Significant differences in the posterior cingulate, precuneus and ventromedial PFC emerged only in run 3, and were evident in both linear and exponential learning contrasts (p < 0.05, whole brain corrected and also following non-parametric tests using threshold-free cluster enhacement). We note that the behavioural expression of the learning effect of the central goal-relevant sequence both in manual RTs and response accuracy followed a different time course to the learning effects found in the brain responses. The former did not significantly increase across the training runs, while learning effects on brain response were most evident in the third training run. It is worth noting that sequence learning effects may reflect a combination of response-response, stimulus-response and response-stimulus bindings (Ziessler & Nattkemper, 2001; Ziessler, 1998), with each having a different time course; however this was not addressed in the present neuroimaging study.

Following prior reports that putative mnemonic regions (e.g., hippocampus) show increased temporal correlation with V1 during memory recall (Bosch et al., 2014) and implicit statistical learning of visible stimuli (Turk-Browne et al., 2010), we conducted similar analyses considering the posterior precuneus cluster that was observed during learning, as this region is also a key node of the memory network (Ranganath & Ritchey, 2012). This region has connections to higher level visual areas (Margulies et al., 2009) and thus has access to V1 via feedback projections.

The psychophysiological interaction (PPI) analysis used a 6 mm seed mask centered on the posterior precuneus area identified in the learning contrast (MNI 0 -48 12) at the cluster peak in the S<R contrast in run 3 reported above. This was used to define the seed region’s time course for the PPI. Given our a priori interest in primary visual cortex, training effects across participants were assessed within a standardized occipital mask as target of the PPI.

We found a cluster of activity in V1 peaking at the intracalcarine sulcus (MNI: 8 -92 0, Z=3.34) showing linear and exponential increases in temporal correlation with posterior precuneus across the training runs on structured relative to the pseudorandom sequences (see figure 4). This result was observed in the comparison between first pseudorandom sequence block and the structure sequence immediately preceding it. This suggests that learning a representation of the structured sequence involves the coupling between putative memory substrates and V1. For the position of anatomically defined V1 in the calcarine, see (Horton & Hoyt, 1991); also, the activity in the intracalcarine cortex and its vicinity coincides with probabilistic V1.

**Figure 4:**
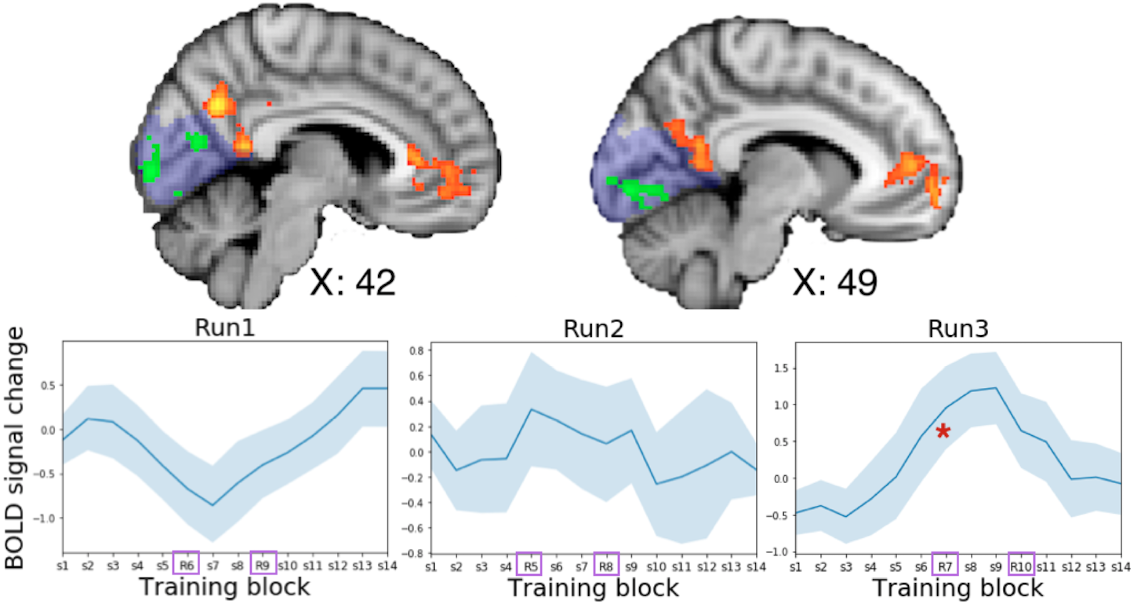
**Top:** Brain regions in the precuneus and ventromedial prefrontal cortex exhibited linear changes in BOLD signal for the structured<pseudorandom sequence contrast across the three training runs (in yellow, Z > 2.3, p < 0.05, whole-brain corrected). The green areas in V1 indicate the results of the PPI analysis from the posterior precuneus cluster (Z > 2.3, p < 0.05, corrected for the occipital mask in blue shading). **Bottom:** Time course of the responses in the posterior precuneus cluster during the training phase as a function of the structured (S) or pseudorandom (R) status of the sequences. The red * illustrates significant BOLD signal change between structured and pseudorandom sequences in run 3.

Right and left V1 were mapped using independent retinotopy and functional localiser scans of each participant (see Methods). On the basis that learning effects (S<R) for the central structured sequence were expressed in the last training run, it was predicted that learning of the peripheral sequences would be also evident in run 3.

Accordingly, the BOLD signal difference between structured and pseudo-random sequences (S<R) in right V1 in the last training period (run 3) was significantly different from chance (Figure 5; t(16)=2.480, p=0.0246, two-tailed paired t-test). Also, we conducted a repeated measures ANOVA with run (1, 2, 3) as factor to test for linear and quadratic trends in the learning effect (S<R for the left sequence) across the runs. There was a significant quadratic trend in the learning effect for right V1 showing that learning effects were expressed in run 3 (F(1,16)=6.32, p<0.023, Figure 5). The linear trend was not significant (F(1,16)=1.39, p<0.25). This pattern of results is consistent with the non-linear learning effects observed for the central sequence. These results were consistent across a wide range of V1 masks. In particular, we initially tested our hypothesis with masks that comprised the 10 most responsive voxels to the location of the left sequence, as shown by the combination of retinotopy and the functional localiser, and then verified that the result held with masks containing 5, 20 and 40 most responsive voxels. Figure 5 illustrates the representative results from the 10 most responsive voxels.

**Figure 5:**
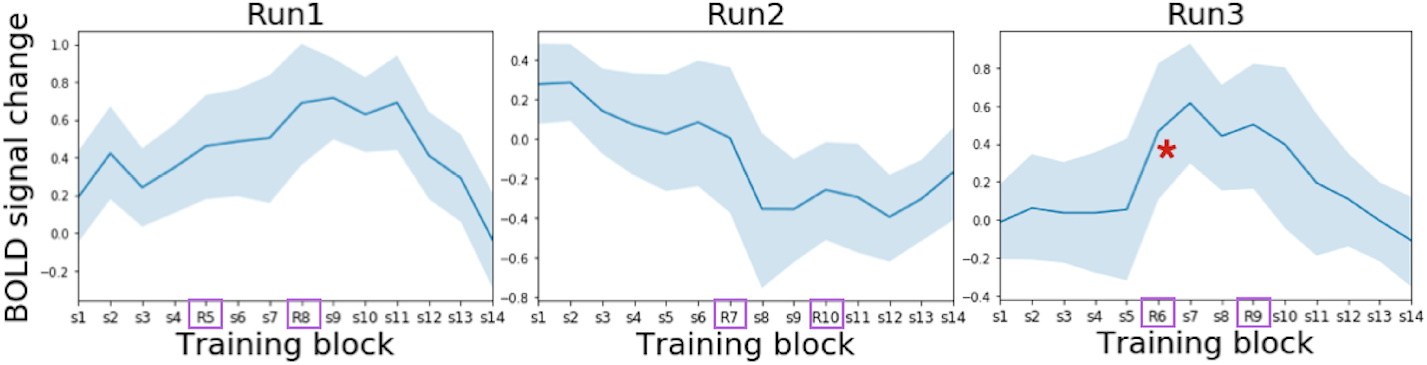
Right V1 responses as a function of the structured (S) or pseudorandom (R) status of the sequences on each of the three training runs. The red * illustrates significant BOLD signal change between structured and pseudorandom sequences in run 3 (see text).

For left V1, there was no evidence of learning the structure of the deterministic discontinuous associations of the structured sequence in the predicted last training run (t(16)=0.66, p=0.518). A direct comparison of the S<R parameter estimates between left and right V1 indicated that the magnitude of the learning effect was higher in right compared to left V1 (t = 2.532, p=0.022, two-tailed paired t-test). Neither the linear nor the quadratic learning trends across runs were observed in the case of left V1 (F(1,16)=.422, p<0.525; (F(1,16)=.829, p<0.376).

Figure 6 below illustrates these results.

**Figure 6:**
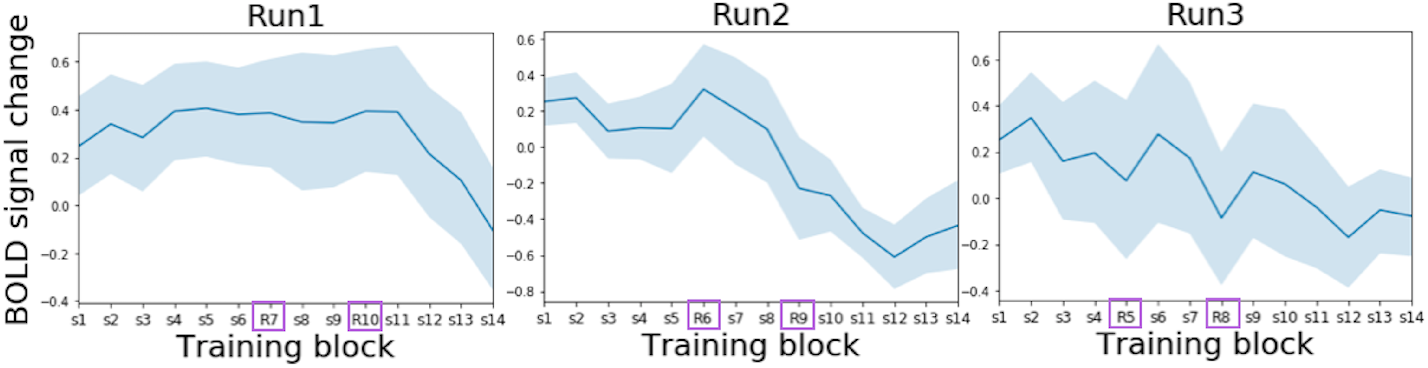
Left V1 responses as a function of the structured (S) or pseudorandom (R) status of the sequences on each of the three training runs. There was no evidence of BOLD signal differences between structured and pseudorandom sequences in any of the runs (see text).

Taken together, these results suggest that the left sequence, but not the right, was encoded in right V1 responses during learning. Additional analysis were conducted to assess whether learning-related activity changes across runs for these peripheral sequences also occurred in similar brain regions that were involved in learning the central sequence. We found that an area in the posterior cingulate at MNI 0 -62 32 showed training-related changes in the S<R BOLD response associated with the left visual sequence (p<0.05, whole-brain corrected), but not the right visual sequence. This area in the posterior cingulate overlapped with that associated with learning the central sequence.

### 3.4 Relationship between BOLD responses during learning and subsequent memory performance for the central sequence

It might be argued that the S<R contrast for the central goal-relevant sequences compares an easier with a more difficult task condition. learning related activity was found in precuneus and vmPFC, which are key nodes of the default mode network that can de-activate at higher levels of task load (Raichle et al., 2001). Here we report results involving correlation analyses between across run learning-related activity changes (S<R) in each of the three different clusters of the network (posterior precuneus, posterior cingulate and ventromedial prefrontal cortex -vmPFC) and subsequent memory performance (i.e., the behavioural assays of newly acquired sequence-specific knowledge). We reasoned that significant correlations would support the view that precuneus and vmPFC activity is, in fact, related to sequence-specific learning rather than task difficulty.

There was a significant correlation between the learning related activity changes in the posterior cingulate and subsequent type-1 memory performance (r=-0.668, p=0.003) and a similar trend was found for type-2 memory performance (r=-0.48, p=0.052). Learning related activity in the precuneus also correlated with type-1 memory performance (r=-0.63, p=0.006) but not type-2 memory performance (r=-0.35, p=0.22). Finally, learning related activity in vmPFC marginally correlated with type-1 memory performance r=-0.47, p=0.054) but no trend was found with type-2 memory performance r=-0.25, p=0.34).Similar analyses were carried out considering the cluster in intracalcarine cortex (V1) that showed increases connectivity with the precuneus during learning. Precuneus-V1 coupling did not correlate with type-1 memory behaviour (r=0.01, p=0.94) and only a marginal negative correlation trend between precuneus-V1 coupling and type-2 memory performance (r=-0.41, p=0.096).

These correlations between posterior cingulate/precuneus and memory performance were negative, meaning that a lower S<R BOLD signal change during training correlated with higher recognition performance. Similar pattern of results have been described previously, namely for memory-related BOLD responses in prefrontal (Grady, McIntosh, & Craik, 2005) and precuneus during memory encoding (Miller et al., 2008). While the specific mechanism that may underlie this association can not be addressed by the present study, these correlations between memory performance and BOLD responses favour an interpretation of the activity in posterior cingulate and precuneus related to the learning of the central sequence rather than stemming from differences in task difficulty between structured and pseudorandom blocks. Also we note here that task difficulty can be operationalised in terms of its effects on performance measures such as reaction time (RT) and/or accuracy. Behavioural analyses showed no indication that the different in performance between structured and pseudorandom blocks was higher in the last training run, while S<R BOLD signal change in posterior cingulate/precuneus and vmPFC significantly increased with training and was higher in the last training run. Therefore, it seems difficult to argue that activity in these areas was associated with changes in performance difficulty.

## 4 Discussion

The results indicate that V1 operates as part of a network involving the precuneus during the learning of goal-relevant visual sequences involving discontinuous associations, which are within the focus of spatial attention. The precuneus, posterior cingulate and ventromedial prefrontal cortex have been identified as part of a memory network (Ranganath & Ritchey, 2012) that can be modulated by the temporal structure of goal-relevant memoranda (Hasson, Chen, & Honey, 2015), with higher-order regions exhibiting sensitivity to increasingly longer temporal structures, ranging from tens of seconds (e.g., written sentences) in more posterior areas to minutes (e.g., a movie).

This cluster of learning related activity in the posterior cingulate/precuneus also extended into the retrosplenial cortex. This area is activated by spatial cognition tasks (Epstein, 2008), episodic memory demands (Vann, Aggleton, & Maguire, 2009) and it has been also heavily implicated in both spatial and object-based forms of contextual/predictive processing and associative learning in scene perception, alongside the ventromedial prefrontal cortex (Bar & Aminoff, 2003; Bar, 2004). V1 coupling with posterior precuneus cortex during repeated exposure to the structured sequences is consonant with a mnemonic feedback signal from higher-order brain regions involved in contextual/predictive processing (Rao & Ballard, 1999; Alink, Schwiedrzik, Kohler, Singer, & Muckli, 2010).

In addition to the contribution of V1 to encoding the central attended sequence, the results revealed that retinotopic V1 areas can also support learning of the statistical regularities associated with goal-irrelevant items presented at non-attended spatial positions. Note that performance in the central attention task based on 4-color discrimination had to be carried out at a fast pace and performance was not at ceiling (i.e., around 0.8 proportion correct). This means that the central task was challenging enough to keep the focus of the observers’ attention engaged, and thus observers were very unlikely to spare attentional resources to process the peripheral 12-element sequences. This is also indicated by the fact that observers performed at chance in the direct tests of subsequent recognition memory for the peripheral sequences. However, the indirect RT-based recognition measures showed differences between old and new sequences. This suggests that participants may have acquired sequence-specific knowledge for the peripheral sequences that could expressed outside the original conditions of training, although it is likely that this knowledge remained implicit given that observers could not discriminate between old and new peripheral sequences using objective and subjective reports.

The present results go beyond a recent demonstration in which V1 was associated with learning of a single second-order conditional sequence (Rosenthal et al., 2016); this study could not determine whether V1 can support multiple co-ocurring sequences and did not address whether learning occurred in retinotopic areas. Furthermore, the learning effects in V1 in Rosenthal et al. were found under conditions in which the elements of the sequence were goal-relevant for a secondary counting test. Thus, the present results show for the first time that V1 can support learning of both goal-relevant and -irrelevant visual sequences presented at retinotopic areas.

The results also indicate that spatial relational encoding is not necessary for the concurrent learning second-order conditional sequences in V1, in keeping with the study of Gavornik and Bear (2014), which used a simpler and single visual sequence. Rosenthal et al.’s (2016) study of non-conscious visual sequence learning employed a single second-order conditional visuospatial sequence specified at four locations. In the present study each of the three second-order conditional sequence were presented at one location. Hence, sequence learning in Rosenthal et al. were characterized by spatial relational encoding, while object-based relational encoding defined the present study. This may be relevant to the presence of basal ganglia activity in Rosenthal et al. and its absence in the present study, since here there is no need for allocentric/egocentric coding of spatial relations, which may be relevant to drive sequence learning effects in the basal ganglia (Kermadi & Joseph, 1995).

Retinotopic activity in right V1 was associated with learning the SOC sequence presented in the left visual field. However, there was no evidence of analogous effects in left V1. It is possible that visual competition amongst the three concurrent sequences reduced overall the processing capacity of left and right V1 areas to encode the multiple co-ocurring sequences, but there is no immediate reason to explain the preferential role of right V1 activity to support sequence-specific learning alongside a null sensitivity in left V1. Prior work has suggested the existence of lateralised perceptual biases (Railo, Tallus, & Hamalainen, 2011; Nicholls, Mattingley, & Bradshaw, 2005) in which perceptual processing is biased towards the left side of space. If such biases were operating here, then this could explain the preferential engagement of right V1 for processing the left sequence. Another possibility to explain the significant learning effect in right V1 and its apparent absence in left V1 may be found by considering a model of visuospatial hemispheric specialization (Kosslyn, 1987). This model proposes that the right hemisphere supports object-oriented processing, which here it would include the serial dependencies between sequence orientations. On the other hand, the model proposed by Kosslyn (1987) argues that processing in the left hemisphere is more general and less object-oriented. In line with this model, memory-based fMRI studies showed that encoding-related activity during study of visual objects in the right fusiform were associated with later remembering of precise object-based relations, while left fusiform activity during encoding predicted subsequent, non-specific recognition (Garoff, Slotnick, & Schacter, 2005).

The learning effects reported in this study are in keeping with recent reports that V1 can be subject to experience-dependent changes that support higher-order mnemonic processes (Rosenthal et al., 2016; Ekman, Kok, & de Lange, 2017; Cooke et al., 2015; Gavornik & Bear, 2014). Gavornik and Bear (2014) reported response increases in V1 activity following the presentation of simple visual sequences that were previously learned. This is in contrast with the activity reduction in V1 seen for the learned sequences in the present study. Notwithstanding, the present results are consistent with our prior study based on learning a single masked visuospatial SOC sequence (Rosenthal et al., 2016) and also with prior sequence learning studies based on visible SOC sequences of stimuli presented at four locations (Schendan et al., 2003; Albouy et al., 2008). It is also largely consistent with human neuroimaging studies of memory in which old<new contrasts associated with recognition memory, cued recall and repetition priming are all related with reduced BOLD responses to ’old’ relative to ’new’ items (Okada, Vilberg, & Rugg, 2012). The sequences of orientation gratings used by Gavornik and Bear (2014) involved simple first-order associations of four elements in length measured across several days of training. Hence, experience-dependent plasticity changes following these conditions are likely to reflect the output of post-learning consolidation mechanisms rather than rapid conjunctive learning of 12-element discontinuous associations (i.e., separated by time) acquired under conditions involving three discrete retinotopic locations and a manipulation involving task relevance. In addition, direct comparisons across species and between the outcome from chronic recording of local field potentials restricted to mouse V1 and human fMRI are challenging.

Our findings suggest that V1 processing is flexible enough to be deployed even when attention is constrained by visual competition and task-relevance factors. It is possible that sequence learning effects are supported by local activity changes in V1 activity. We also acknowledge the possibility that the pseudorandom sequences captured attention due to the violation of the predictability. Accordingly V1 activity could have been mediated by attentional modulation. However, there is at least one factor which argues against this possibility. In particular, the central task imposed a high attentional load which likely reduced the ability of the pseudorandom sequences to capture attention. Previous studies have shown that a central attentional load impairs detection of even simple visual stimuli (Lavie, 2005; Lavie, Beck, & Konstantinou, 2014). Note also there were no overall differences in task demands between structured and pseudorandom sequences, the stimuli that composed both sequence types were equally familiar - the presence of discontinous SOC associations was the only attribute that distinguished structured from the pseudorandom sequences-, and, given that there was evidence of retrieval success of the peripheral sequences evinced by differences in manual RTs, we favour an interpretation where the S<R signal change in V1 reflected the (en)coding of sequence-specific structural properties of the SOC which seems needed in order to trigger a ’novelty’ response to the R sequence.

Top-down processes related to attention (Kastner, Pinsk, De Weerd, Desimone, & Ungerleider, 1999) and predictive processing (Kok, Jehee, & De Lange, 2012) are known to modulate V1 (Watanabe et al., 2011). However, that V1 is capable of encoding complex knowledge at goal-irrelevant peripheral locations is consistent with a recent observation that V1 can automatically ‘preplay’ an expected sequence of events, even when attention resources are constrained (Ekman et al., 2017). Further research is needed to establish the extent to which complex learning effects in V1 are mediated by local plasticity changes, and the extent of the potential contribution of top-down feedback processes. However, based on the observation that learning modulated retinotopic V1 for peripheral items that were goal-irrelevant, whereas attentional resources were exhausted by a central task load, we tentatively suggest that V1 local plasticity changes are likely to play a role.

Finally, it is worth noting that the mnemonic feedback signals we have observed in V1 may come from different laminae than feedforward signals; experiments at higher resolution will be needed to make this determination.

## 5 Acknowledgements

D.S. acknowledges support from the Spanish Ministry of Economy and Competitiveness (MINECO), through the ’Severo Ochoa’ Programme for Centres/Units of Excellence in R&D (SEV-2015-490) and project grants PSI2016-76443-P from MINECO and PI-2017-25 from the Basque Government.

